# LADL: Light-activated dynamic looping for endogenous gene expression control

**DOI:** 10.1101/349340

**Authors:** Mayuri Rege, Ji Hun Kim, Jacqueline Valeri, Margaret C. Dunagin, Aryeh Metzger, Wanfeng Gong, Jonathan A. Beagan, Arjun Raj, Jennifer E. Phillips-Cremins

## Abstract

Mammalian genomes are folded into tens of thousands of long-range looping interactions^1,2^. The cause and effect relationship between looping and genome function is poorly understood, and the extent to which chromatin loops are dynamic on short time scales remains a fundamental unanswered question. Currently available strategies for loop engineering involve synthetic transcription factors tethered to dCas9^3,4^ or zinc fingers^5,6^, which are constitutively expressed^5,6^ or induced on long time scales by the presence of a small molecule^3^. Here we report a new class of 3-D optoepigenetic tools for the directed rearrangement of 3-D chromatin looping on short time scales using blue light. We create synthetic architectural proteins by fusing the CIBN protein subunit from *Arabidopsis thaliana* with enzymatically dead Cas9 (dCas9). We target our light-activated dynamic looping system (LADL) to two genomic anchors with CRISPR guide RNAs and engineer their spatial co-localization via light-induced heterodimerization of the cryptochrome 2 (CRY2) protein with dCas9-CIBN. We apply LADL to redirect a stretch enhancer (SE) away from its endogenous *Klf4* target gene and to the *Zfp462* promoter. Looping changes occur as early as four hours after light induction. Using single molecule RNA FISH, we observe a LADL-induced increase in the total nascent *Zfp462* transcripts and the number of *Zfp462* alleles expressing simultaneously per cell. Moreover, LADL also increased synchronous *Sox2* expression after reinforcement of a known *Sox2*-SE looping interaction. LADL facilitates loop synchronization across a large population of cells without exogenous chemical cofactors and can enable future efforts to engineer reversible and oscillatory looping on short time scales.

## Main

The development of tools to manipulate 3D genome folding on demand with spatiotemporal precision is of critical importance toward advancing studies in basic science, regenerative medicine, metabolic engineering and synthetic biology. Our light-activated dynamic looping system (LADL) was modularly designed with four key components (**Figure 1a**). First, we designed a synthetic architectural protein consisting of enzymatically inactive Cas9 (dCas9) tethered to a truncated CIBN protein derived from the CIB1 protein from *Arabidopsis thaliana* (**Figure 1b**). Second, we recruited the LADL Anchor (dCas9-CIBN) to two genomic target sites with sequence-specific CRISPR guide RNAs (gRNAs) (**Figure 1c**). Two gRNAs were designed per anchoring genomic target site. Third, the PHR domain of the CRY2 protein from *Arabidopsis thaliana* was used as the bridging protein due to its well-established ability to heterodimerize with CIBN in response to blue light on millisecond time scales in mammalian cells^7,8^ (**Figure 1c**). Finally, the inducing agent is blue light of wavelength 470nm^9^. It is well-established that blue light illumination causes CIBN-CRY2 heterodimerization^7,8^ and CRY2 oligomerization^7,10,11^, thus we hypothesized that LADL would spatially connect the two anchoring genomic fragments via a light-induced dCas9-CIBN and Cry2 bridge (**Figure 1a**). In the dark condition, dCas9-CIBN binds to the two genomic anchors while CRY2 remains soluble. Plasmid design is detailed in the Supplementary Methods (**Supplementary Figures 1-17, Supplementary Tables 1-6)**. Thus, LADL is designed as a modular, four-component architectural protein system, which connects looping interactions in response to light via facile design of new sequence-specific gRNAs.

**Figure 1.**
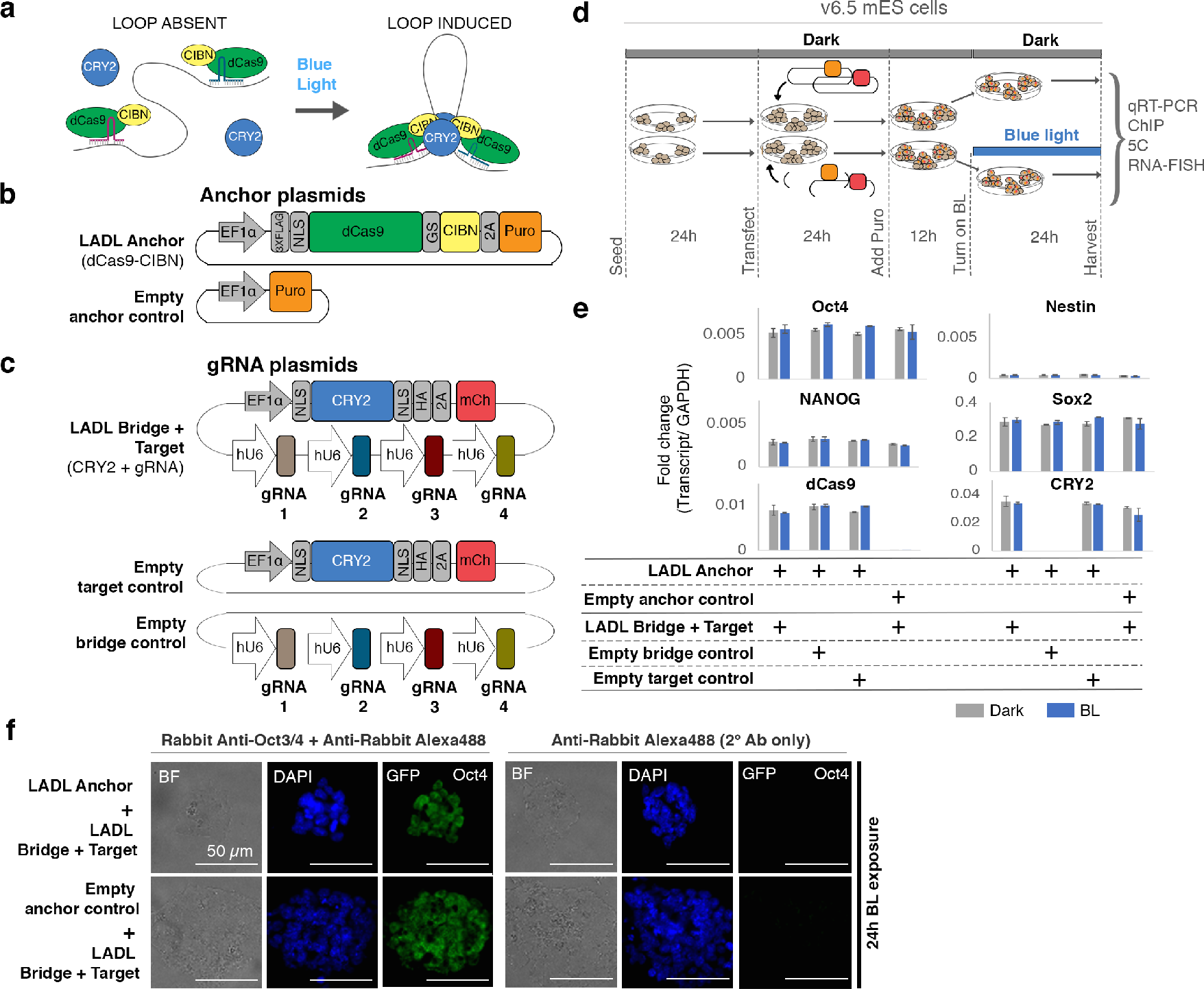
Concept, design and implementation of the light-activated dynamic looping (LADL) system. **(a)** LADL schematic. In the dark, the LADL anchoring protein dCas9-CIBN is recruited to two specific genomic fragments using guide RNAs (gRNAs). Cry2 is soluble and does not specifically associate with chromatin. Upon illumination with blue light, Cry2 homodimerizes and Cry2 and dCas9-CIBN heterodimerize, forming a bridge that connects the two gRNA-bound genomic fragments and loops out the intervening DNA. **(b-c)** Schematic of the constructs encoding the **(b)** puromycin-selectable LADL Anchor and Empty anchor control plasmids and **(c)** the LADL Bridge + Target, the Empty target control and the Empty bridge control plasmids. **(d)** Schematic timeline of seeding, transfection, puromycin selection and light illumination of v6.5 mouse embryonic stem cells. **(e)** qRT-PCR analysis of *Oct4*, *Nestin*, *Nanog*, *Sox2*, *dCas9*, and *CRY2* transcript levels in co-transfected mouse embryonic stem cells after 36 hours post-puromycin selection. Two biological replicates each. ANOVA followed by Tukey’s HSD test was used to compare differences in the means. No significant differences were found. Two biological replicates each. Error bars represent standard deviation. **(f)** Immunofluorescence staining for Oct4 in mouse embryonic stem cells co-transfected with the indicated plasmids.
Scale bars are 50 μm and under 40X objective magnification.

To determine the conditions in which blue light would induce a spatial chromatin contact, we first employed 24 hours of blue light exposure (**Figure 1d**). A light box was built to achieve 5 mW/cm2 intensity and 1-second pulses at 0.067 Hz as previously reported to optimally facilitate CRY2-CIBN heterodimerization^9,12^ (detailed in **Supplementary Methods**). We confirmed that these light exposure conditions successfully induced Cry2 oligomerization **(Supplementary Figure 14).** Mouse embryonic stem (ES) cells were co-transfected to achieve one of four conditions: (1) LADL (Anchor (dCas9-CIBN) and Bridge+Target (CRY2+sgRNA) plasmids), (2) Empty anchor control (Empty anchor and Bridge+Target (CRY2+gRNA) plasmids), (3) Empty bridge control (Anchor (dCas9-CIBN) and Empty bridge (gRNA only) plasmids), or (4) No target control (Anchor (dCas9-CIBN) and Empty target (Cry2 only) plasmids). Overall plasmid mass and ratios were adjusted to optimize transfection efficiency (**Supplementary Figure 15**).

Transfected cells were exposed to 24 hours of blue light or dark after puromycin selection (**Figure 1d**). ES cell densities were similar across conditions and exhibited morphology characteristic of the v6.5 feeder-dependent clone after passage onto gelatin (**Supplementary Figure 16**). All three conditions showed equivalently high expression of pluripotency markers *Oct4*, *Nanog*, *Sox2*, and low levels of *Nestin*, suggesting that the pluripotent, self-renewing ES cell state was not compromised by transfection and light induction (**Figure 1e-f, Supplementary Table 7).** *dCas9-CIBN* and *CRY2* transcripts were strongly expressed across all conditions transfected with vectors encoding the transgenes **(Figure 1e)**. Moreover, equivalent levels were seen in dark and blue light exposure, thus ruling out artifacts caused by differential transgene levels between conditions. Together, these results demonstrate that the two plasmids encoding our 4-component synthetic architectural protein system were equivalently expressed and have minimal negative impact on ES cell morphology, viability, and pluripotent properties.

We next set out to choose the genomic context for our LADL engineered loop testing. The *Klf4* and *Zfp462* genes are highly and lowly expressed in pluripotent ES cells, respectively, and are under the control of distal enhancer elements (**Figure 2**). As previously reported^13^, *Zfp462* forms long-range interactions with up to four independent putative enhancers (E1, E2, E3, E4) marked by positive enrichment of the histone modification H3K27ac (**Figure 2a**). *Klf4* forms a ~70 kb-sized long-range interaction with a large putative stretch enhancer (SE)^13,14^. We reasoned that we could test LADL’s looping ability by relying on a ‘Redirect and Reinforce’ strategy in which we spatially redirected the *Klf4* SE away from *Klf4* and to the *Zfp462* promoter. To avoid disrupting endogenous transcription factors and architectural proteins, we designed LADL gRNAs directly adjacent to, but not overlapping, the H3K27ac ChIP-seq and ATAC-seq signal marking the *Klf4* SE and the *Zfp462* promoter (**Figure 2a-c**, blue and red gRNA markers respectively, **Supplementary Table 7**). We used chromatin immunoprecipitation followed by quantitative PCR (ChIP-qPCR) to confirm recruitment of the LADL system to the specifically targeted genomic locations (**Figure 2d-f, Supplementary Table 8**). Using an anti-FLAG antibody, we demonstrated strong enrichment of FLAG-tagged dCas9-CIBN in the dark at both the *Zfp462* promoter (**Figure 2e**) and the *Klf4* SE (**Figure 2f**), but not a non-specific genomic region (**Figure 2d**). Importantly, this enrichment was not observed when the LADL Anchor was absent (Empty anchor control). Thus, the LADL Anchor can be effectively targeted to genomic loci adjacent to accessible chromatin using two gRNAs.

We next set out to determine if a spatial contact was induced by LADL in response to blue light. We hypothesized that an engineered long-range contact between our two targeted genomic fragments might alter dCas9-CIBN ChIP-qPCR signal due to indirect immunoprecipitation from the distal, spatially proximal fragment (**Figure 2g**). Indeed, we found that the intensity of dCas9-CIBN ChIP signal is altered after blue light illumination, increasing more than 2-fold at the *Zfp462* promoter and slightly decreasing at the *Klf4* SE (**Figure 2e-f**). We then directly assessed higher-order chromatin architecture with Chromosome-Conformation-Capture-Carbon-Copy (5C)^13,15,16^ (**Supplementary Tables 9-10**). We generated a high-resolution map of long-range interactions for all genomic fragments in a ~3.5 Mb region around the *Klf4* and *Zfp462* genes in the conditions of (1) LADL (Anchor + Bridge + Target) in 4 and 24 hours of blue light, (2) LADL (Anchor + Bridge + Target) in 4 and 24 hours of dark and (3) Empty target control (Anchor+Bridge only) in 4 and 24 hours of dark (**Figure 3a-b, Supplementary Figure 17**). Upon blue light illumination, a new long-range contact is gained between the SE and *Zfp462* in mouse ES cells transfected with LADL vectors (**Figure 3c–e, Supplementary Figure 18**). The engineered loop is specific to the LADL+blue light condition and not present in LADL+dark or Empty target+dark controls (**Figure 3c-e, Supplementary Figure 18**). We reproduced the *de novo Zfp462-Klf4* SE loop in LADL-transfected ES cells after 24 hours of blue light illumination at a lower intensity of 1.5 mW/cm^2^ (**Supplementary Figure 19**). Together, these results demonstrate that LADL can form a new long-range interaction between two genomic fragments in a blue light-dependent manner. We noticed a specific disruption to chromatin architecture that occurred in the process of redirecting the *Klf4* SE to *Zfp462*. In wild type mouse ES cells, the *Klf4* gene forms a strong looping interaction with its target stretch enhancer; this loop is detectable in both LADL (Anchor + Bridge + Target) in dark condition and the Empty target control (Anchor + Bridge only) in dark condition (**Figure 3f-g, Supplementary Figure 20**). Consistent with our objective to redirect the *Klf4* SE away from its endogenous target gene, the punctate looping interaction between the SE and *Klf4* is markedly reduced in LADL transfected ES cells upon blue light illumination (**Figure 3f-g, Supplementary Figure 20**). Moreover, *Zfp462* creates a network of long-range interactions with multiple pluripotency-specific enhancers^13,14^. The enhancers also interact with each other, but this hub of enhancer-enhancer and enhancer-*Zfp462* interactions was undisturbed across all conditions **(Supplementary Figure 21).** These data demonstrate that the *Klf4* SE can be redirected across a population of cells to *Zfp462*, consequently breaking down endogenous *Klf4*-enhancer interactions, whereas endogenous *Zfp462*-enhancer interactions remain unaffected (**Figure 3h**).

**Figure 2.**
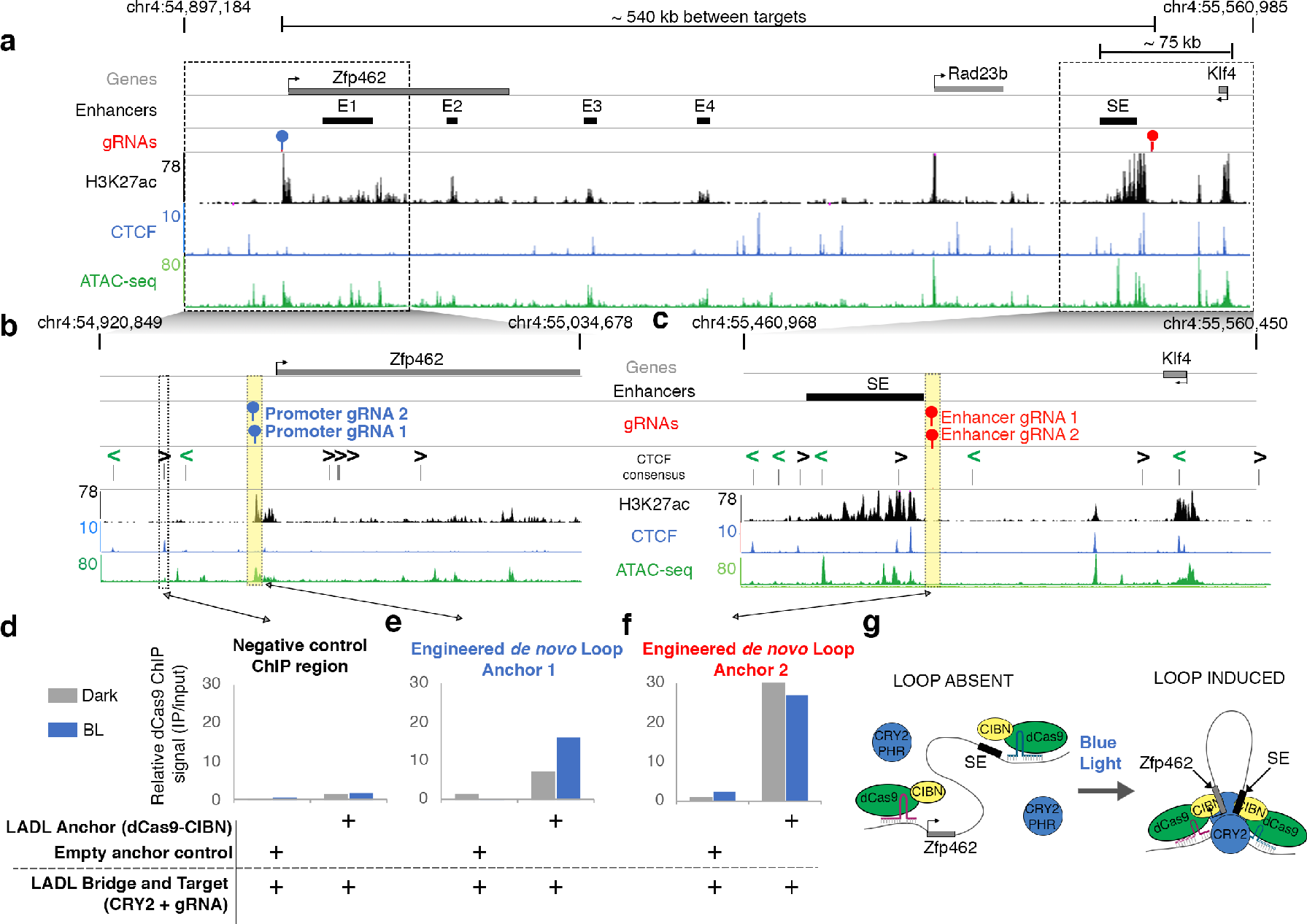
Chromatin binding of LADL Anchor (dCas9-CIBN) at the engineered sites increases after blue light exposure. **(a)** Genome browser tracks overlaid at the *Zfp462* and *Klf4* genes and their target enhancers (chr4:54,897,184-55,560,985; mm9 reference genome). SE, the *Klf4* super enhancer. E1, E2, E3, E4, the *Zfp462* enhancers. **(b-c)** Zoomed-in genome browser tracks at (**b)** the *Zfp462* promoter and **(c)** adjacent to the SE. **(d-f)** ChIP-qPCR data for the **(d)** negative control chromatin site, **(e)** engineered gRNAs at the *Zfp462* promoter and **(f)** engineered gRNAs at the *Klf4* SE in co-transfected mouse embryonic stem cells after 24 hours of dark or blue light exposure. **(g)** Model illustrating findings from the ChIP-qPCR data.

**Figure 3.**
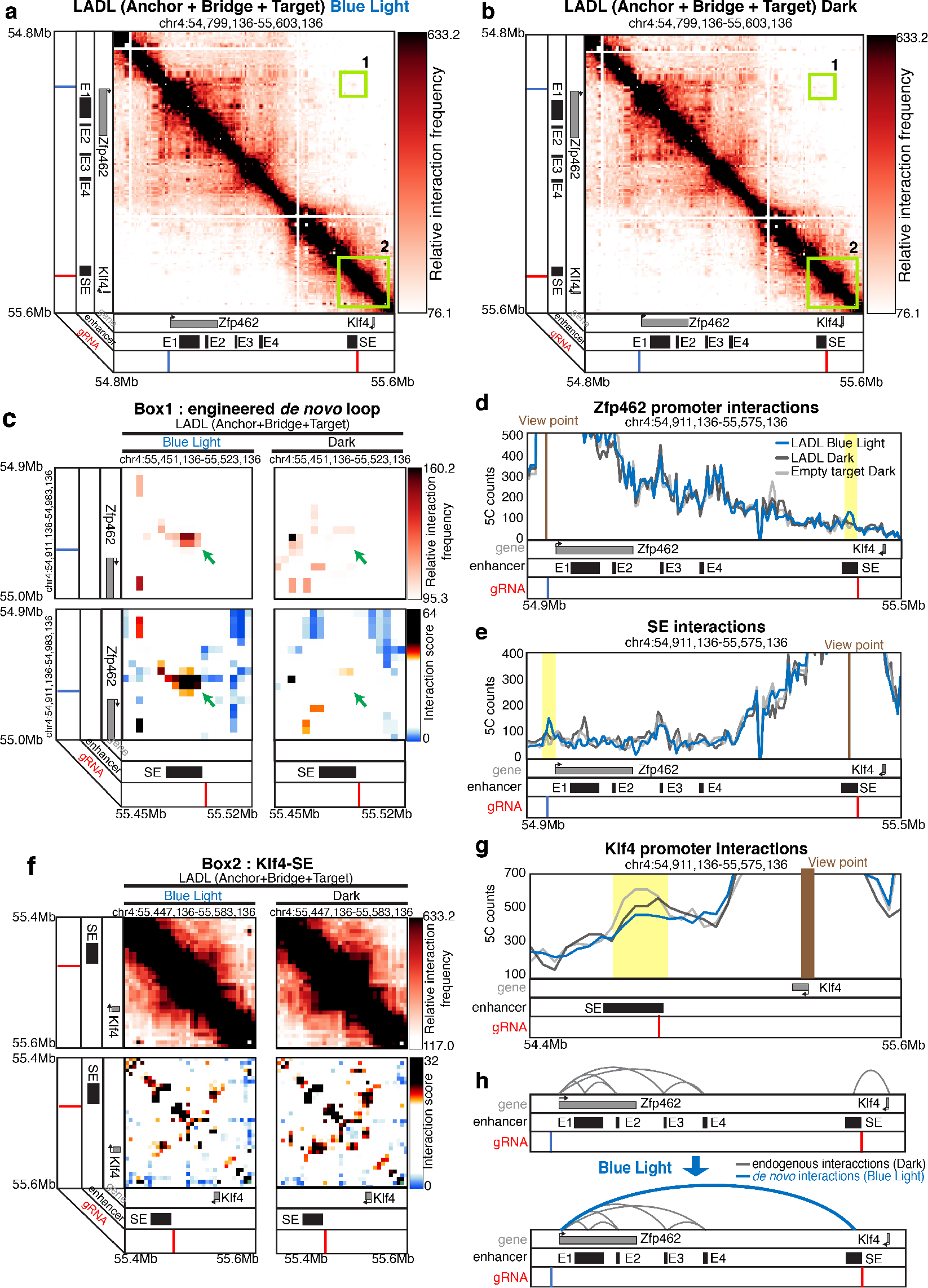
LADL redirects and reinforces a long-range interaction between a stretch enhancer and a new target gene upon blue light illumination. **(a-b)** Heatmap of long-range interactions around an ~800 kb genomic region encompassing the *Klf4* and *Zfp462* genes. SE, the *Klf4* super enhancer. E1, E2, E3, E4, the *Zfp462* enhancers. Mouse embryonic stem cells were co-transfected with LADL (Anchor + Bridge + Target) plasmids and then exposed to 24 hours **(a)** blue light illumination or **(b)** dark. Box 1, the target de novo engineered loop between the pluripotency-specific *Klf4* stretch enhancer and the *Zfp462* transcription start site. Box 2, *Klf4* interaction with its upstream, pluripotency-specific stretch enhancer. The Empty target control (Anchor+Bridge only) plus dark is shown in **Supplementary Figure 17**. **(c, f)** Zoomed-in heatmaps of **(c)** Box 1 or **(f)** Box 2. Top, relative interaction frequency 5C signal. Bottom, distance-corrected interaction score 5C signal. The Empty target control (Anchor+Bridge only) plus dark is shown in **Supplementary Figures 18+20**. **(d, e, g)** Classic 4C interaction frequency plots from the viewpoint of **(d)** the *Zfp462* promoter targeted gRNA, **(e)** the stretch enhancer targeted gRNA, or **(g)** the *Klf4* promoter. **(h)** Model of looping interaction reconfiguration in response to LADL and blue light illumination.

To gain insight into the time scale upon which the LADL-induced loops are formed, we mapped chromatin architecture in LADL engineered ES cells with 5C after varying the time scale of blue light exposure (**Figure 4a**). We observed that the *de novo* engineered loop between *Zfp462* and the *Klf4* SE occurred after as little as 4 hours blue light illumination (**Figure 4b-d, Supplementary Figure 22**). Importantly, the LADL-engineered loop exhibited similar interaction strength in 4 hour vs. 24 hour blue light exposure (**Figure 4e**). Consistent with this finding, the *Klf4*-SE interaction was reduced in LADL-engineered ES cells after both 4 and 24 hours blue light compared to the LADL+dark and Empty target+dark conditions (**Supplementary Figure 23**). Together, these results indicate that LADL can enable the formation of long-range looping interactions on-demand in as early as 4 hours after application of the induction stimulus. Chemical induction of looping is reported to occur on the time scale of 24 hours or more^3^, thus LADL provides a significant advance in shortening the time scale of loop induction.

**Figure 4.**
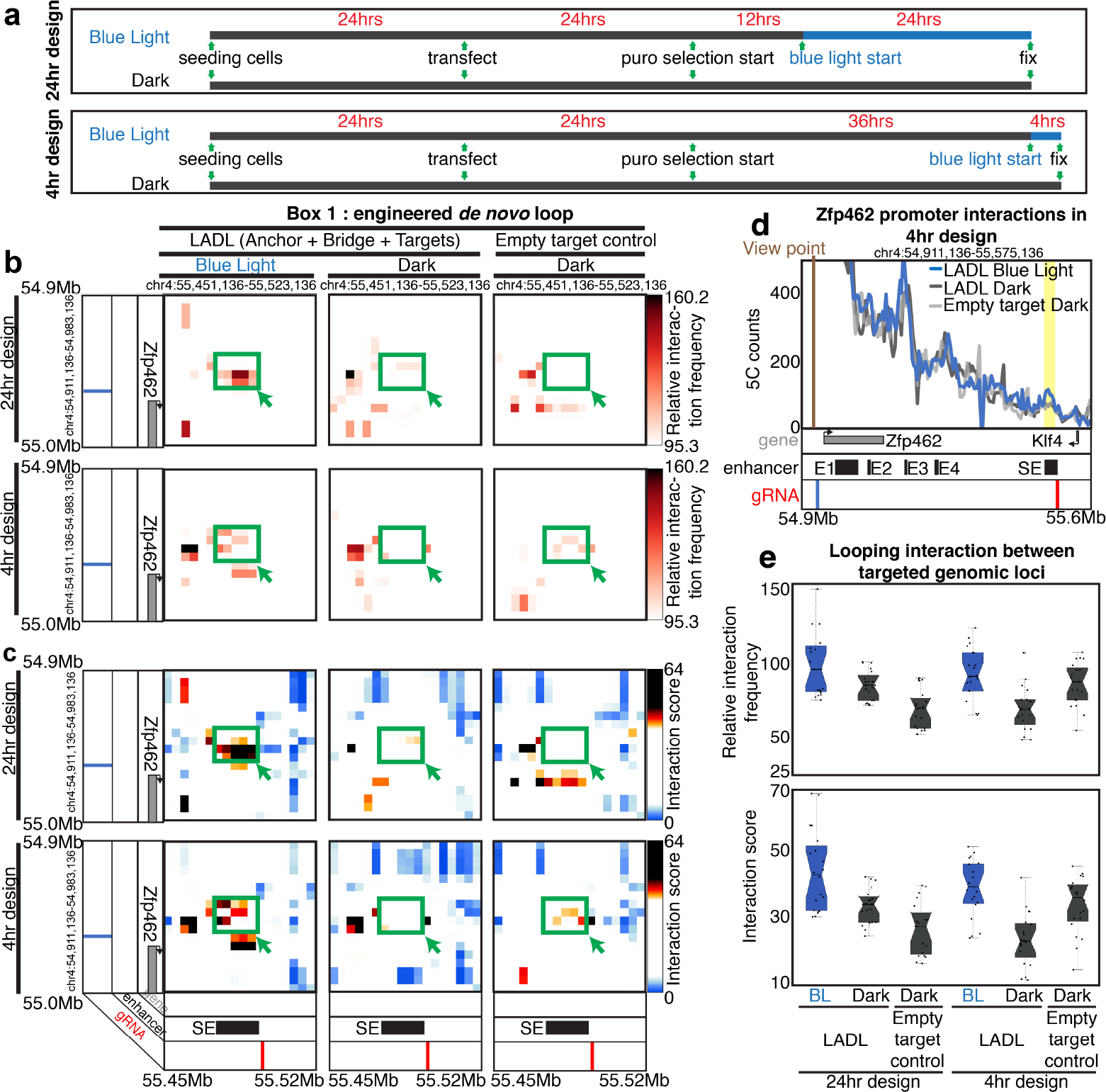
The LADL-induced long-range interaction occurs as early as 4 hours after blue light illumination. **(a)** Schematic of LADL experimental design with 4 and 24 hours of blue light illumination. Mouse embryonic stem cells were co-transfected with LADL (Anchor + Bridge + Target) plasmids or Empty target control (Anchor + Bridge only) plasmids. At the time of harvesting, the cells have undergone puromycin selection for at least 36 hours in either 4 or 24 hour blue light exposure. **(b-c)** Zoomed in heatmaps corresponding to Box 1 from Figure 3a-b for **(b)** Relative interaction frequency 5C signal and **(c)** Distance-corrected interaction score 5C signal. **(d)** Classic 4C interaction frequency plots from 4 hour blue light illumination experiment from the viewpoint of *Zfp462* promoter targeted gRNA. **(e)** Boxplots of both relative interaction frequencies (top) and distance-corrected interaction scores (bottom) in the *de novo* engineered loop pixels (green box in **b-c**) for all 6 conditions from 4 and 24 hour blue light exposure designs.

We next set out to understand the functional effects of LADL-engineered looping interactions on gene expression. We performed single molecule RNA FISH to assess *Zfp462* and *Klf4* expression changes on a single cell basis after 36 hours puromycin treatment to select only LADL-transfected cells and 24 hours blue light illumination (**Figure 5a, Supplementary Table 13**). The mean number of total *Zfp462* mRNA transcripts per cell (53.47; 95% CI: 48.54 < μ_*Zfp462*_LADL+BL_ < 58.40) was significantly higher in LADL+BL compared to LADL+dark (43.68; 95% CI: 38.24 < μ_*Zfp462*_LADL+dark_ < 49.11), Empty target control+dark (40.14; 95% CI: 36.69 < μ_*Zfp462*_Empty target control+BL_ < 43.49), or Empty bridge control+dark (40.33; 95% CI: 37.24 < μ_*Zfp462*_Empty bridge control+BL_ < 43.41) (**Figure 5b**). Moreover, the mean number of nascent *Zfp462* transcripts per cell (3.40; 95% CI: 3.02 < μ_*Zfp462*_LADL+BL_ < 3.78) was significantly higher in LADL+BL compared to LADL+dark (2.72; 95% CI: 1.96 < μ_*Zfp462*_LADL+dark_ < 2.59), Empty target control+dark (2.57; 95% CI: 2.33 < μ_*Zfp462*_Empty target control+BL_ < 2.81), or Empty bridge control+dark (2.63; 95% CI: 2.34 < μ_*Zfp462*_Empty bridge control+BL_ < 2.93) (**Figure 5c**). Thus, these results indicate that the functional consequence of forced looping between the *Klf4* SE and the *Zfp462* promoter is an increase in total mRNA and nascent *Zfp462* transcripts per cell (**Figure 5e**).

We reasoned that *Klf4* expression might be affected by LADL, given the *Klf4*-SE interaction is reduced as the SE is redirected to *Zfp462.* Indeed, total mRNA levels of *Klf4* per cell were shifted downward (71.63; 95% CI: 62.35 < μ_*Klf4*_LADL+BL_ < 80.91) in LADL+BL compared to LADL+dark (85.34; 95% CI: 72.63 < μ_*Klf4*_LADL+dark_ < 98.05), Empty target control+dark (78.01; 95% CI: 65.99 < μ_*Klf4*_Empty target control+BL_ < 90.03), or Empty bridge control+dark (102.08; 95% CI: 88.39 < μ_*Klf4*_Empty bridge control+BL_ < 115.77) (**Figure 5b**). Moreover, total levels of nascent *Klf4* transcripts per cell were also shifted downward (1.09; 95% CI: 0.81 <μ_*Klf4*_LADL+BL_ < 1.37) in LADL+BL compared to LADL+dark (1.30; 95% CI: 1.0 < μ_*Klf4*_LADL+dark_ < 1.60), Empty target control+dark (1.12; 95% CI: 0.85 < μ_*Klf4*_Empty target control+BL_ < 1.39), or Empty bridge control+dark (1.79; 95% CI: 1.46 < μ_*Klf4*_Empty bridge control+BL_ < 2.13) (**Figure 5c**). In both cases, a statistically significant reduction in *Klf4* was achieved when comparing LADL+BL to only the Empty target control+dark condition. These data are consistent with the hypothesis that the redirection of *Klf4*’s SE to a different target gene might reduce its expression in parallel with the reduction of the *Klf4*-SE loop (**Figure 5e**).

**Figure 5.**
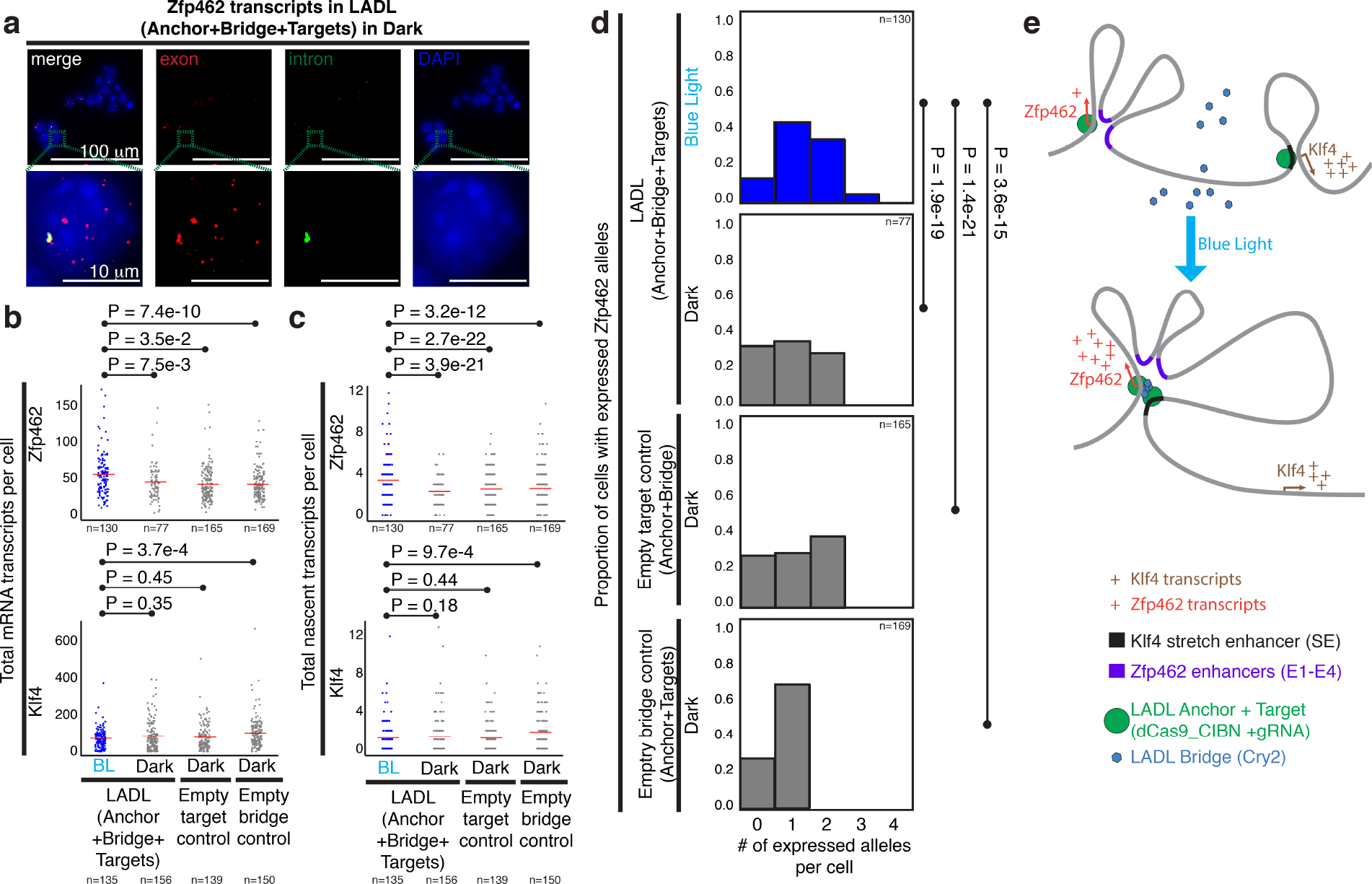
Functional effect of the LADL-engineered de novo loop on endogenous gene expression. **(a)** Full field (top row) and zoomed microscopic images (bottom row) from single molecule RNA fluorescence *in situ* hybridization (FISH) analysis of LADL-engineered mouse embryonic stem cells in dark. After blue light illumination at 5 mW/cm^2^ for 24 hours, *Zfp462* or *Klf4* transcripts in LADL-engineered mouse embryonic stem cells and in three other controls (LADL+dark, Empty target control+dark, Empty bridge control+dark) were hybridized with 32 exon-and 32 intron-specific fluorescence-labeled oligonucleotides before acquiring images for quantitative analysis (detailed in Supplementary Methods). **(b-c)** Strip charts representing **(b)** the total number of mRNA transcripts per cell and **(c)** the total nascent transcripts per cell for *Zfp462* (upper row) and *Klf4* (lower row). Red bars indicate means of each condition. P-values computed using the non-parametric Mann Whitney U test. **(d)** Histograms represent the proportion of cells with a specific number of actively expressing *Zfp462* alleles. P-values computed using the non-parametric Mann Whitney U test. **(e)** Schematic diagram illustrating the link between LADL-engineered de novo loop formation and changes in endogenous gene expression.

We also hypothesized that synchronized looping might lead to an increase in the number of cells within a population that simultaneously express a particular gene. Using single molecule RNA FISH, we observed that the number of alleles per cell expressing *Zfp462* was significantly increased in LADL+BL (1.32; 95% CI: 1.19 < μ_*Zfp462*_LADL+dark_ < 1.46) compared to LADL+dark (0.97; 95% CI: 0.79 < μ_*Zfp462*_LADL+dark_ < 1.16), Empty target control+dark (1.12; 95% CI: 0.99 < μ_*Zfp462*_Empty target control+BL_ < 1.24), or Empty bridge control+dark (1.02; 95% CI: 0.91 < μ_*Zfp462*_Empty bridge control+BL_ < 1.14) (**Figure 5d**). The extent to which loops are dynamic and heterogeneous across individual cells in a population is unknown. However, our data suggest that LADL effectively redirects the *Klf4* SE away from its *Klf4* target to the *Zfp462* promoter and synchronizes the engineered loop to coordinate an increase in *Zfp462* transcription across a population of cells (**Figure 5e, Supplementary Figure 24**).

In addition to the *de novo* redirect-reinforce looping strategy, we wanted to assess whether LADL could reinforce an existing loop to strengthen and synchronize gene expression. A hallmark of the pluripotent stem cell state is the presence of a strong looping interaction between the Sox2 gene and its target SE. To test the capability of LADL to reinforce the *Sox2*-SE interaction, we screened a series of gRNA strategies. First, we pursued a ‘Hijack and Reinforce’ strategy in which we recruited gRNAs directly to the CTCF binding sites anchoring the base of the *Sox2* promoter and SE loop (**Supplementary Figure 25, 28a-f**). Second, we formulated an alternative ‘Hijack and Reinforce’ strategy in which we targeted CTCF sites located between the two regulatory elements (**Supplementary Figure 26, 28a-f**). Finally, we tested a ‘Reinforce’ strategy in which we target so-called desert loci devoid of chromatin marks between *Sox2* and its SE (**Supplementary Figure 27, 28a-f**).

To assay the effect of the different gRNA strategies on gene expression, we used the well-characterized Sox2-eGFP induced pluripotent stem (iPS) cell line that expresses the Sox2-eGFP protein from the endogenous Sox2 promoter ^17^. We transfected Sox2-eGFP iPS cells with either the LADL constructs or the Empty target control under BL or dark conditions and quantified the GFP intensity per transfected cell (**Supplementary Figure 28g-h**). Of the three gRNA strategies, only “Hijack & Reinforce Strategy 1” increased the Sox2-eGFP levels significantly on exposure to 5 mW/cm2 of BL for 24h (**Supplementary Figure 28h**). Moreover, the fraction of cells transfected with the “Hijack & Reinforce Strategy 1” that exhibit a high intensity of GFP is significantly higher with LADL compared to dark or Empty target controls (**Supplementary Figure 28i**). These results demonstrate that LADL can induce the upregulation of endogenous gene expression due to light activated looping across different genomic contexts.

Overall, we present LADL as a new set of synthetic architectural proteins that form inducible long-range looping interactions in response to light. The use of our new ‘3D optoepigenetic tools’ to engineer chromatin topology will be useful because they: (1) facilitate loop activation and reversibility on rapid time scales, (2) enable the previously unachievable ability to oscillate spatial contacts, (3) overcome signal/noise issues in population-based genomics assays by synchronizing chromatin topology across a large population of cells in response to blue light illumination and (4) are scalable *in vivo* and will allow for clinical applications that involve spatial targeting of specific neurons in the brain. To our knowledge, LADL represents a novel synthetic architectural looping system controlled by light, opening up future opportunities for spatial targeting of specific cells *in vivo* for dynamic looping and control of gene expression on short time scales.

## Acknowledgements

We thank members of the Cremins lab for helpful discussions. Jennifer E. Phillips-Cremins is a New York Stem Cell Foundation – Robertson Investigator and an Alfred P. Sloan Foundation Fellow. This research was supported by The New York Stem Cell Foundation (J.E.P.C), the Alfred P. Sloan Foundation (J.E.P.C), the NIH Director’s New Innovator Award from the National Institute of Mental Health (1DP2MH11024701; J.E.P.C), a 4D Nucleome Common Fund grant (1U01HL12999801; J.E.P.C), and a joint NSF-NIGMS grant to support research at the interface of the biological and mathematical sciences (1562665; J.E.P.C).

## Data Access

All raw data, code and analysis files will be freely and publicly available at publication.

## Author contributions

J.E.P.C., M.R., J.V., A.M. conceptualized the system. M.R., J.H.K., J.V., W.G. designed and performed the experiments. M.D. and A.R. designed and conducted RNA FISH experiments. M.D. A.R. and J.E.P.C. analyzed FISH data. J.H.K., J.A.B., J.E.P.C. performed the 5C data analysis. J.H.K performed Sox2-eGFP quantitative microscopy analysis. J.E.P.C. wrote the manuscript with help from all authors.

## Draft Competing interests

A patent disclosure has been filed relating to this work.

